# Accumulation of copy-back viral genomes during respiratory syncytial virus infection is preceded by diversification of the copy-back viral genome population followed by selection

**DOI:** 10.1101/2022.05.25.493350

**Authors:** Sébastien A. Felt, Emna Achouri, Sydney R. Faber, Carolina B. López

## Abstract

RNA viruses generate non-standard viral genomes during their replication, including viral genomes of the copy-back (cbVG) type that cannot replicate in the absence of a standard virus. cbVGs play a crucial role in shaping virus infection outcomes due to their ability to interfere with virus replication and induce strong immune responses. However, despite their critical role during infection, the principles that drive the selection and evolution of cbVGs within a virus population are poorly understood. As cbVGs are dependent on the virus replication machinery to be generated and replicated, we hypothesized that host factors that affect virus replication exert selective pressure on cbVGs and drive their evolution within a virus population. To test this hypothesis, we used respiratory syncytial virus (RSV) as model and took an experimental evolution approach by serially passaging RSV in immune competent A549 control and immune deficient A549 STAT1 KO cells which allow higher levels of virus replication. As predicted, we observed that virus populations accumulated higher amounts of cbVGs in the more permissive A549 STAT1 KO cells over time but, unexpectedly, the predominant cbVG species after passages in the two conditions were different. While A549 STAT1 KO cells accumulated relatively short cbVGs, A549 control cells mainly contained cbVGs of much longer predicted size that have not been described previously. These long cbVGs were predominant at first in both cell lines *in vitro* and the predominant ones observed in samples from RSV infected patients. Although sustained high replication levels are associated with cbVG generation and accumulation, our data show that sustained high levels of virus replication are critical for cbVG population diversification, a process that preceded the generation of shorter cbVGs that selectively accumulated over time. Taken together, we show that selection and evolution of cbVGs within a virus population is shaped by how resistant or permissive a host is to RSV.

## 1. Introduction

The ability of RNA viruses to rapidly evolve is a major concern for public health due to the potential emergence of virus variants with enhanced pathogenicity (Carrasco-Hernandez et al., 2017). It is, therefore, of critical importance to understand the principles that drive virus evolution. An RNA virus consists of communities of standard viral genomes and non-standard viral genomes, such as copy-back viral genomes (cbVGs), that are generated during virus replication (Vignuzzi and Lopez, 2019). cbVGs are generated when the polymerase detaches from the template strand at a break point and resumes elongation at a downstream rejoin point creating a complementary end to the 5’-end of the nascent genome (Kolakofsky, 1976; Lazzarini et al., 1981; Vignuzzi and Lopez, 2019). cbVGs are usually shorter than standard viral genomes and their flanking trailer promoters have a higher affinity for the viral polymerase than the leader promoter present at the 3’ end of the standard viral genome, thereby providing a replication advantage to cbVGs (Li and Pattnaik, 1997; Rao and Huang, 1982). This advantage in replication can lead to the accumulation (significant increase) of shorter cbVGs within virus populations. cbVGs depend on standard viral genomes to be generated and replicated. Many cbVGs are packaged, released from the infected cell, and can infect new host cells affecting the interaction of the virus with the host (Lazzarini et al., 1981; Vignuzzi and Lopez, 2019). cbVGs can inhibit standard viral genome replication by competing for the virus replication machinery or by inducing innate immune signaling pathways (Henle and Henle, 1943; Strahle et al., 2006; Sun et al., 2015; Tapia et al., 2013). Despite the critical role of cbVGs in shaping the infection outcome, most RNA virus evolution studies only consider standard viral genomes. As a result of this, the principles that drive the selection and evolution of cbVGs within a virus population are poorly understood.

As cbVGs depend on the virus replication machinery to be generated and amplified, we hypothesized that host factors that affect virus replication exert selective pressure on cbVGs and drive their evolution within a virus population. To test this hypothesis, we used the nonsegmented negative sense RNA virus respiratory syncytial virus (RSV) as a model because RSV generates many cbVGs during infection *in vitro* and *in vivo* (Collins et al., 2013; Felt et al., 2021; Sun et al., 2015; Treuhaft and Beem, 1982). RSV cbVGs stimulate MAVS signaling during infection, which leads to the production of antiviral type I and III interferons (IFNs) (Sun et al., 2019). This antiviral response is dependent on cbVGs triggering the intracellular viral sensors RIG-I-like receptors, as has been shown for many non-segmented negative sense RNA viruses (Baum et al., 2010; Ho et al., 2016; Linder et al., 2021; Mura et al., 2017; Runge et al., 2014; Strahle et al., 2007; Xu et al., 2015; Yount et al., 2008). The expression of type I and III IFNs leads to the activation of STAT1, which is followed by the production of hundreds of proteins that interfere with virus replication (Mesev et al., 2019). Interestingly, recently it has been shown that viruses can activate STAT1 even earlier independently of IFNs through a RIG-I/MAVS/Syk pathway further establishing STAT1 as an important antiviral protein (Liu et al., 2021). *In vivo*, RSV can replicate to higher levels in STAT1 knockout mice compared to wildtype mice (Durbin et al., 2002; Hashimoto et al., 2005; Stier et al., 2017). Based on these studies, we decided to test if STAT1 exerts selective pressure on RSV cbVGs and drive their evolution. Specifically, we took an experimental evolution approach and serially passaged RSV using a fixed-volume or a fixed-MOI in immune competent A549 control cells or in immune deficient A549 STAT1 KO cells. As RSV also generates cbVGs in infected patients (Felt et al., 2021), we also analyzed cbVGs in nasal washes from RSV infected pediatric patients to assess what cbVGs are selected for during natural infections.

To detect cbVGs in RNA-seq data, we used an updated version of our previously published Viral Opensource DVG Key Algorithm (VODKA) (Sun et al., 2019), called VODKA2, which identifies cbVG reads based on their alignment to unique junction sequences that contain a break and rejoin point. Predicted cbVG species sizes can then be calculated based on the break and rejoin points. Our study revealed that the diversity of cbVGs generated is much larger than previously thought. We found that cbVGs of longer predicted sizes that have not been described previously are predominant at first *in vitro* and found predominantly in samples from RSV infected patients, and that sustained high virus replication levels led to cbVG population diversification that preceded the generation and selective accumulation of shorter cbVGs.

## 2. Methodology

### 2.1 Cells and viruses

A549 control cells, A549 STAT1 KO cells and HEp2 cells (HeLa-derived human epithelial cells, ATCC, CCL23) were cultured at 5% CO_2_ and 37°C with Dulbecco’s modified Eagle’s medium supplemented with 10% fetal bovine serum (FBS), 1 mM sodium pyruvate, 2 mM L-Glutamine, and 50 mg/ml gentamicin. The A549 CRISPR cell lines were kindly provided by Dr. Susan Weiss (University of Pennsylvania) and have been previously characterized (Whelan et al., 2019; Xu et al., 2017). All cell lines were treated with mycoplasma removal agent (MP Biomedicals) and routinely tested for mycoplasma before use. RSV A2 was obtained from ATCC (#VR-1540) and amplified in Hep2 cells at a low MOI of 0.01 to generate RSV stocks with low cbVG contents (Sun and Lopez, 2016). We used these stocks as parental (P0) virus stocks for our experiments.

### 2.2 Passaging experiments

In fixed-volume passaging experiments, for the first passage, A549 control and STAT1 KO cells were infected at an MOI of 10 with an RSV stock with low cbVG content (P0) to focus mainly on *de* novo generation of cbVGs. We chose this MOI as higher MOIs favor the accumulation of non-standard viral genomes, including cbVGs, in virus populations (Huang, 1973; Kingsbury et al., 1970; Stampfer et al., 1971; Sun and Lopez, 2016; Thompson and Yin, 2010; Von Magnus, 1954; von, 1951; Welch et al., 2020). Two days post infection (before cells start dying significantly (Collins et al., 2013)), cells were collected and centrifuged for 5 minutes at 280 x g to separate supernatant from cells. Most of the supernatant was transferred to a new tube and kept on ice to preserve viral particles. The remaining supernatant was left to cover the cell pellet to make sure the cell pellet does not dry. As most RSV particles remain attached to cells (Collins et al., 2013), three freeze/thaw (dry ice/ethanol + 37C water bath) cycles were performed on the cell pellet to release virus particles. Cells were vortexed after each freeze/thaw cycle to further increase virus particle release. Cells were then centrifuged for 5 minutes at 280 x g to separate cell debris from supernatant with released viral particles. Supernatant from freeze/thaw cycles was then mixed with supernatant that was kept on ice and 36 ul, which represent ~1/25^th^ of the total mixed supernatant, was then used to infect the next round of fresh cells (passage 2). These procedures and volume were kept consistent throughout the remaining passages. We chose a ratio of ~1/25 as in a previous study it was shown to lead to accumulation of non-standard viral genomes (Williams et al., 2016). At each passage the remaining supernatant was snap frozen in dry ice/ethanol to preserve viral particles for downstream analysis (titration and RNAseq/VODKA2 analysis). It is important to note that, in addition to viral particles, the samples also contain unpackaged viral RNAs.

For the fixed-MOI passaging experiment, A549 control and STAT1 KO cells were initially infected at an MOI of 0.1 or 10 with an RSV stock with low cbVG content (P0). The steps between each passage and preservation of samples were performed such as for the fixed-volume experiment. However, supernatants were tittered in between each passage to maintain the same MOI throughout the 10 passages. For the fixed-MOI of 10 experiment, an MOI of 10 was maintained if titer and volumes allowed it. For the first 6 passages it was possible to use an MOI of 10, however, for passages 7, 8, 9 and 10, the MOIs were reduced to 5, 8, 5 and 5 respectively.

### 2.3 Titration

Supernatants were tittered by TCID_50_ on Hep2 cells to assess the amount of infectious viral particles as previously published (Sun and Lopez, 2016). Briefly, supernatants were serially diluted, added on Hep2 cells, incubated for 4-5 days and after addition of crystal violet the last dilution with CPE was determined to calculate the TCID_50_.

### 2.4 Nasopharyngeal aspirates

Nasopharyngeal aspirates from pediatric patients were obtained from the Children’s Hospital of Philadelphia (CHOP). The CHOP cohort was previously described in detail (Felt et al., 2021). All samples used were banked samples obtained as part of standard testing of patients. Samples were de-identified and sent to our lab for RNA extraction and cbVG detection.

### 2.5 RNASeq

Total RNA was extracted using TRIzol LS (Invitrogen). RNA quality was assessed using an Agilent TapeStation or Bioanalyzer (Agilent Technologies) prior to cDNA library preparation. All samples were prepared using the Illumina Non-Stranded Total RNA Library Prep Kit with Ribo-Zero Gold. The only exceptions were the clinical samples, which were prepared using the Sigma SeqPlex RNA Amplification Kit and the samples of Figure S3, which were prepared using the Illumina TruSeq Stranded Total RNA Library Prep Kit with Ribo-Zero Gold. All samples were run on a NovaSeq 6000 to generate 150 bp, paired-end reads, resulting in ~33–62 million reads per sample. The only exceptions were samples of Figure S3, which were run on an Illumina NextSeq 500 to generate 75 bp, single-end reads, resulting in ~ 45–72 million reads per sample. Only R1 reads were used for analysis as many reads in R2 were duplicates. Average phred quality score of samples ranged between 34.7-36.1. Raw RNAseq data from Figure S3 and Figure S5A were used in previous studies (Sun et al., 2019; Xu et al., 2017).

### 2.6 Viral Opensource DVG Key Algorithm 2 (VODKA2)

After trimming sequencing adapters (lllumina Universal Adapter – AGATCGGAAGAG, https://support.illumina.com/) from the 3’ end of the reads using Cutadapt (DOI:10.14806/ej.17.1.200) and removing all reads aligned to the human genome (GRCh38, GRC website) with Bowtie2 (https://doi.org/10.1038/nmeth.1923), an updated version of VODKA (VODKA2.0 beta, https://github.com/eachouri/VODKA2/tree/main/VODKA2-v2.0b-master), was used to detect cbVG junction reads deriving from the entire viral genome (*in vitro* reference genome KT992094.1 and clinical samples reference genome KC731482.1). We only focused on cbVGs generated from the 5’ end of the viral genome, which are the most described in the literature (Vignuzzi and Lopez, 2019). Briefly, VODKA2.0 beta first runs a Bowtie2 alignment of the reads against a database representing all theoretical cbVGs potentially present in the dataset (KT992094.1: 231,298,470 possibilities and KC731482.1: 231,541,872 possibilities). Reads are removed unless they map across break and rejoin junction sequences with at least 15bp of mapped segment on each side. This is followed by an extra filtering step based on the Basic Local Alignment Search Tool (Blastn v.2.11.0, https://doi.org/10.1016/S0022-2836(05)80360-2) alignment of the predicted break and rejoin junction sequences against the virus reference genome (BLAST options: ‘-word_size 11 - gapopen 5 -gapextend 2 -penalty −3 -reward 2 -evalue 0.001 -perc_identity 0.1’). Only sequences with two alignment ranges reported on opposite strands of the reference genome are considered cbVG reads. Sequences with BLAST alignment positions that are not consistent with the initial Bowtie2 alignment are discarded. An extended report is then generated, including in particular the aggregation of cbVG junction reads into ‘species’, i.e. junction reads leading to cbVGs of exact same theoretical size but with break and rejoin occurring within a range of +/-5 nucleotides. ‘Species’ with only 1 read were removed from analysis. Break and rejoin positions from VODKA2.0 beta were used to calculate the predicted size of cbVGs [(break position-genome size)+(rejoin position-genome size)+2].

### 2.7 Statistical analysis

Statistical analyses were performed as indicated in each figure legend using GraphPad Prism v.9.

## 3. Results

### 3.1 cbVGs accumulate to higher levels after 20 passages in A549 STAT1 KO cells than in A549 control cells

To examine the evolution of cbVG populations and selection of cbVG species, we passaged a fixed volume of supernatant from RSV infected cells twenty times in A549 control cells or A549 STAT1 KO cells **(Figure 1A)**. To assess *de novo* generation of cbVGs we started with a parental virus stock (Passage 0, P0) that contained low levels of cbVGs. We confirmed the low cbVG content by our previously established RT-PCR method and RNAseq/VODKA2 pipeline **(Table 1)** (Sun et al., 2015; Sun et al., 2019). We carried out three experimental evolution replicates (3 lineages) for each cell line and repeated this experiment independently three times. For simplicity in visual representation, one representative repeat is shown throughout. As expected, RSV replication, as measured by titer, was overall higher in A549 STAT1 KO cells than in A549 control cells, at least for the first 14 passages, for all three lineages **(Figure 1B)**. RSV replication dropped to very low levels for several passages in A549 control cells, whereas RSV replication was maintained to high levels throughout the entire experiment in A549 STAT1 KO cells **(Figure 1B).** To assess how cbVGs evolved in these two cell lines, we performed RNASeq followed by analysis using VODKA2 for the detection of cbVGs in P0 and in passage 20. cbVG junction reads were detected in all samples and cbVG content increased significantly more in virus populations passaged in A549 STAT1 KO cells (S1-S3) than in A549 control cells (C1-C3) **(Table 1 and Figure 1C)**. These data demonstrate that an environment that does not allow RSV to maintain high replication levels, such as A549 control cells, impedes cbVG accumulation. However, an environment that allows RSV to maintain high replication levels, such as A549 STAT1 KO cells, promotes cbVG accumulation. This is consistent with the literature as most studies use permissive cells or cells with innate immune defects to generate viral stocks with high cbVG content (Santak et al., 2015; Sun and Lopez, 2016; Tilston-Lunel et al., 2021; Welch et al., 2020).

**Figure 1:**
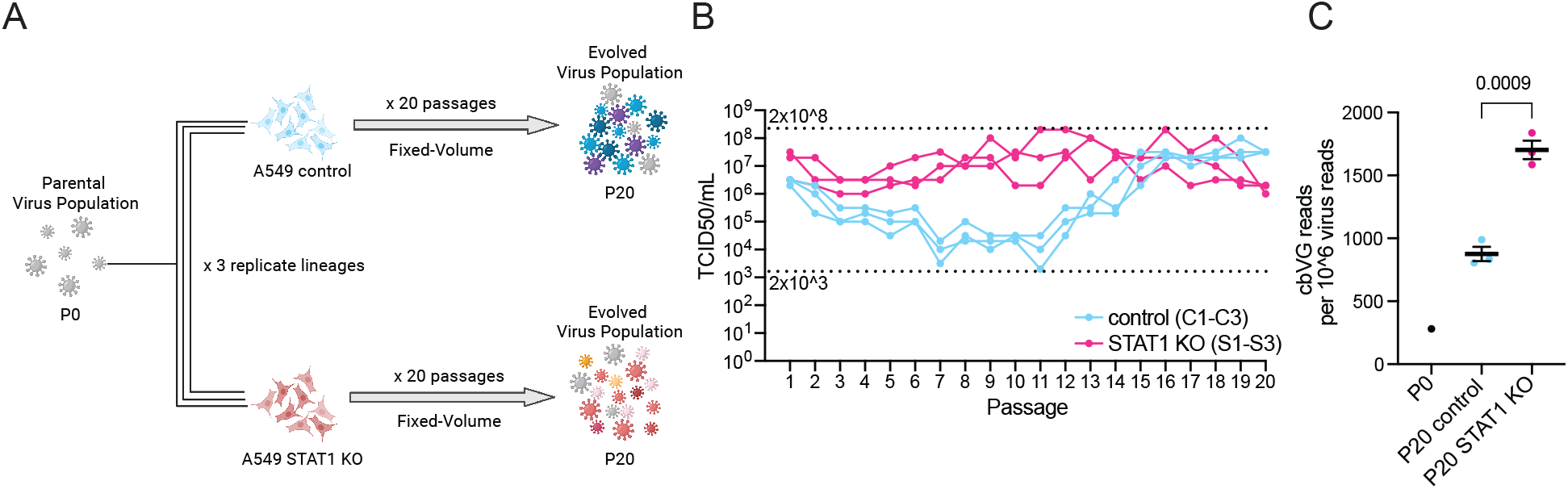
Experimental evolution design and detection of infectious viral particles and cbVGs. (A) Schematic of experimental evolution experiment. (B) TCID_50_ assay was performed on Hep2 cells for all samples. Dashed lines indicate the minimum and maximum TCID_50_/ml values reached in the experiment. (C) cbVG reads detected by VODKA2 were normalized per every 1 million virus reads. Each blue or pink dot represents a lineage. Black dot represents P0. Data are shown as mean ± s.e.m. Significant P value for two-tailed unpaired t test between A549 control and A549 STAT1 KO (n = 3 lineages) is indicated.

**Table 1:**
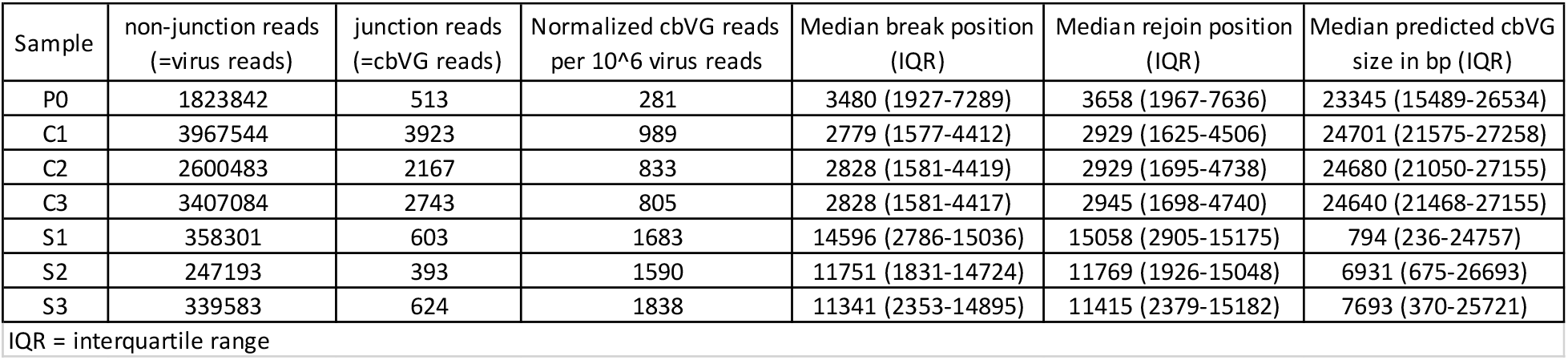
RNAseq/VODKA2 data of parental virus population and passage 20 virus populations.

### 3.2 Short cbVGs accumulate after 20 passages in A549 STAT1 KO cells but not in A549 control cells

We next determined the break (where the polymerase fell off) and rejoin (where the polymerase re-attached) genome position of the cbVGs present in each sample **(Figure S1A)**. As cbVGs unique sequence and size can be predicted by its break/rejoin positions, we used these data to determine if different cbVG species were present in our samples. In previous studies, we set up VODKA to detect cbVGs generated only on the last 3000 bp of the RSV genome as we were only interested in the cbVG species with the most reads (Sun et al., 2019). However, in this study, we assessed the entire cbVG population and thus we set up VODKA2 to identify cbVG junction reads generated through the entire RSV genome.

We observed that more cbVGs in virus populations from A549 STAT1 KO cells had break and rejoin positions in the last 3000bp of the RSV genome (red box) corresponding to cbVG species of predicted shorter size as compared to virus populations from A549 control cells **(Figure 2A)**. This is consistent with our previous study where we showed that RSV populations with high cbVG content had break and rejoin hotspots towards the trailer end of the RSV genome (Sun et al., 2019). However, by running VODKA2 on the entire genome, we discovered novel cbVG species with break and rejoin positions outside the 3000 bp range and many with predicted sizes longer than the genome **(Figure 2A and Figure S1B)**. It is also important to note that most cbVG species detected in P20 were not present in P0. We found that between 90.5% to 95.9% of cbVG species in the six P20 cbVG populations were *de novo* generated rather than being carry-over from P0.

**Figure 2:**
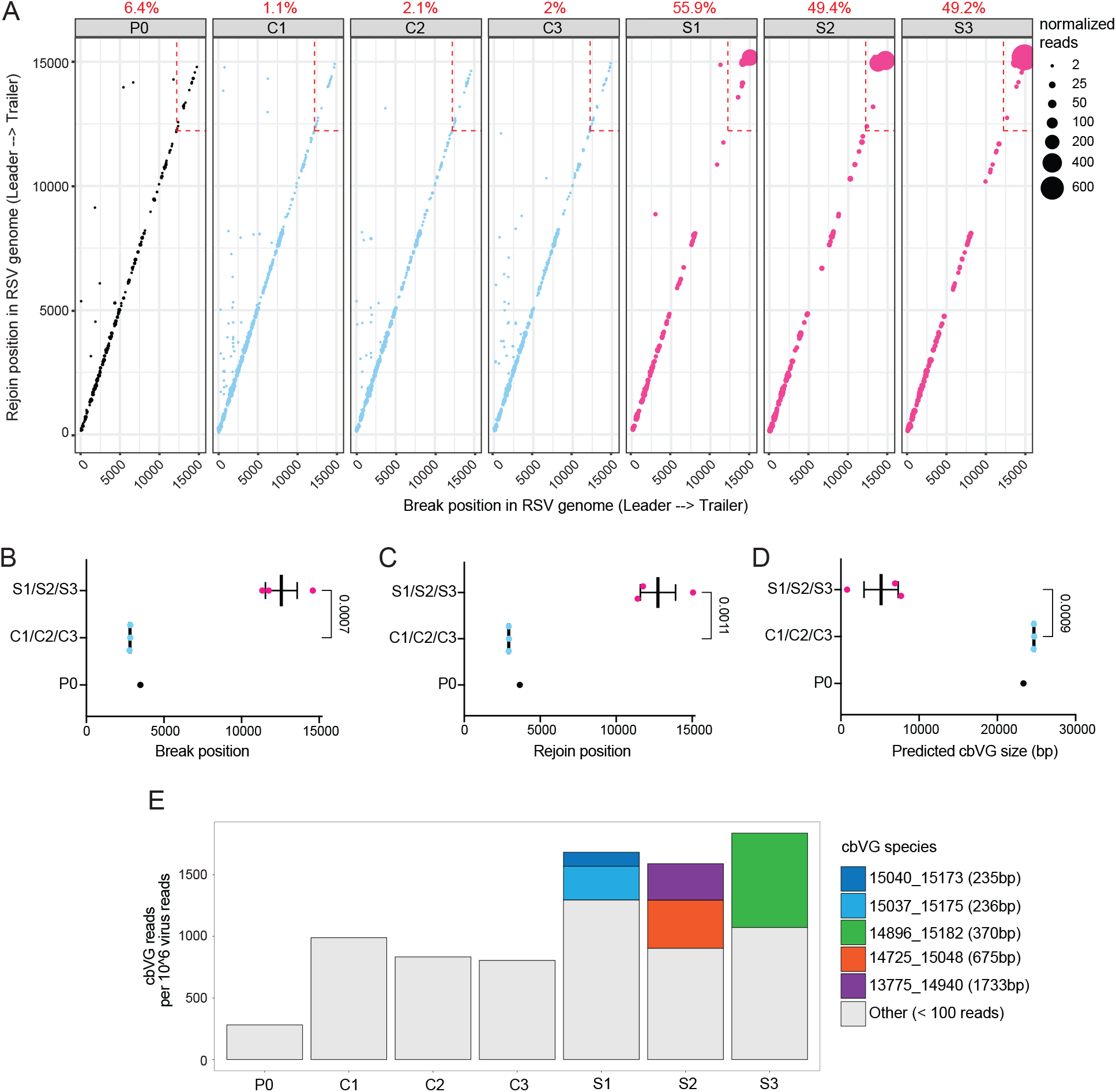
cbVG species in A549 control and A549 STAT1 KO cells at passage 20. (A) Each dot represents a cbVG species and its break/rejoin position. The size of the dots represents the number of normalized reads per 10^6 virus reads. A legend was added as a reference for size. The black dots are cbVGs species from P0, the blue dots are cbVG species from A549 control cells, the pink dots are cbVG species from A549 STAT1 KO cells. The percentage of cbVG reads within the last 3000bp of the RSV genome (red box) are indicated in red above each plot. (B-D) Mean break position, rejoin position and predicted cbVG size are shown in these graphs. Each dot represents the median of a lineage. Data are shown as mean ± s.e.m. Significant P values for two-tailed unpaired t test between A549 control and A549 STAT1 KO (n = 3 lineages) are indicated. (E) Stacked bar graph shows cbVG species with at least 100 reads present in each sample. The legend includes information on break position, rejoin position and predicted cbVG size. cbVG species with less than 100 reads were grouped together as “Other”.

To consider variability and stochasticity in cbVG species generation and accumulation, we also plotted the median break **(Figure 2B)**, rejoin **(Figure 2C)**, and predicted size **(Figure 2D)** for each lineage **(Table 1)**. The median values of the 3 lineages in each condition were similar and the differences observed in break, rejoin, and predicted size in different conditions were statistically different **(Figure 2B-D)**. Finally, we observed that in each A549 STAT1 KO cells lineage (S1-S3) cbVG species of short sizes started to dominate the cbVG population after 20 passages **(Figure 2A and 2E)**. Taken together, our data show that cbVG populations are diverse, that cbVGs are generated across the entire genome and that short cbVG species are selected for in permissive A549 STAT1 KO cells. This selection was not observed in virus populations passaged in more resistant A549 control cells. Instead, long cbVGs remained predominant and no individual longer cbVG species outcompeted the others.

### 3.3 Longer predicted cbVGs are predominant in nasal washes of hospitalized RSV-infected pediatric patients

We detected predominantly cbVG species of longer predicted sizes in virus populations from A549 control cells **(Figure 2A)**. Longer predicted cbVG species were also present in virus populations from A549 STAT1 KO cells, however they were outcompeted by shorter cbVG species **(Figure 2A and 2E)**. In both cell lines, longer predicted cbVGs species had fewer reads, which indicates they do not accumulate like the shorter cbVG species **(Figure 2A)**. To determine whether long cbVGs are of biological significance, we sought to assess if these longer predicted cbVGs could be detected during natural infections. We have previously shown that detection of RSV cbVGs in nasal secretions of infected patients is associated with distinct clinical outcomes. To determine what cbVG species are generated in natural RSV infections, we chose six representative samples from RSV-infected pediatric patients that were previously confirmed to contain cbVGs by our pan-cbVG species RT-PCR approach (Felt et al., 2021). We first extracted RNA from the nasal washes and analyzed them following our RNAseq/VODKA2 pipeline. We observed that cbVGs generated within the last 3000bp of the genome (red box) were almost absent in the samples, and mostly cbVG species of longer predicted size were detected **(Figure 3 and Table 2)**. These results indicate that cbVGs species of a longer predicted length are also generated during natural infections and are the most predominant cbVG species. Overall, although cbVGs of longer predicted size do not accumulate as much as short cbVGs *in vitro* **(Figure 2A and 2E)**, these newly discovered species of cbVGs are predominant *in vivo* **(Figure 3)** and associate with distinct clinical outcomes as reported in our previous study (Felt et al., 2021), emphasizing the importance of taken them in consideration when studying RSV cbVG populations.

**Figure 3:**
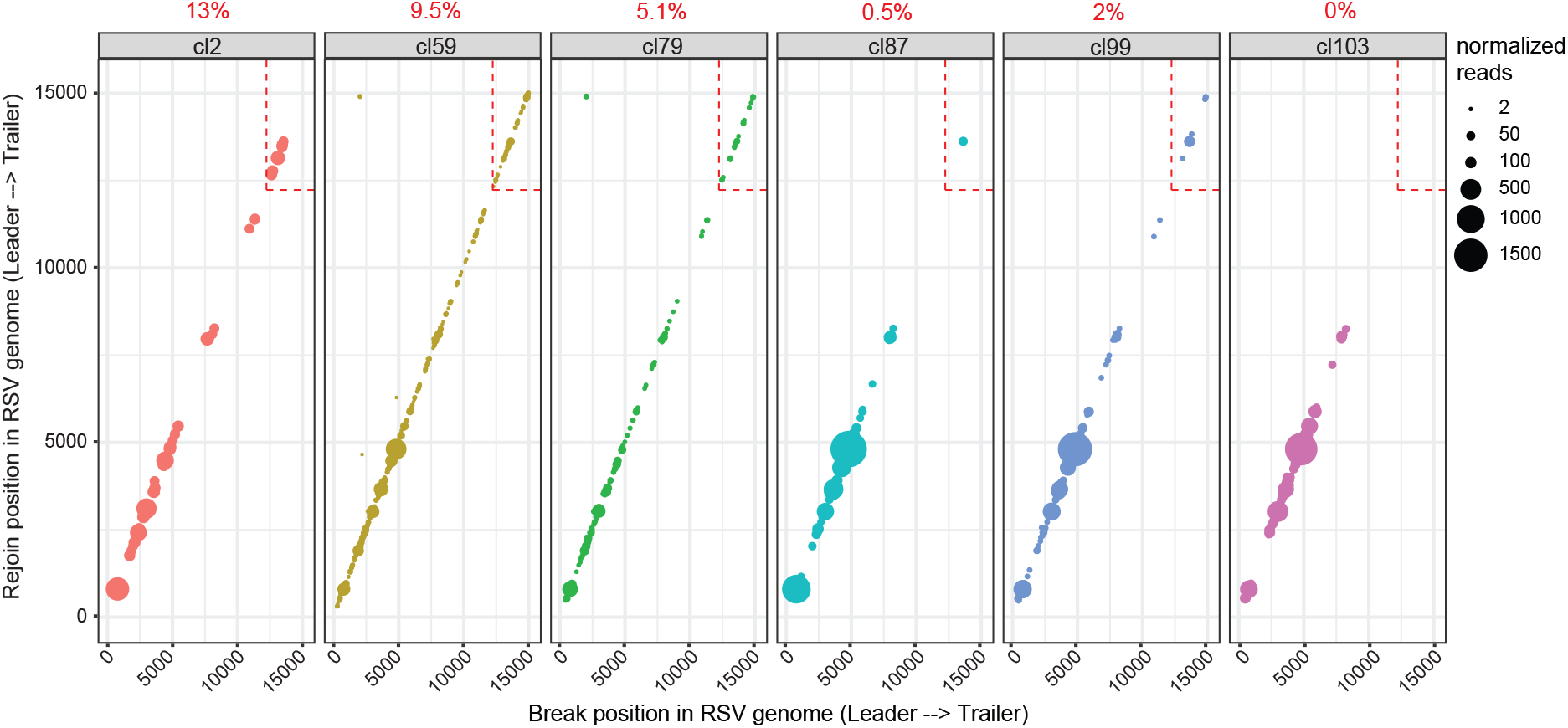
cbVGs in nasal washes of RSV-infected hospitalized pediatric. Each dot represents a cbVG species and its break/rejoin position. The size of the dots represents the number of normalized reads per 10^6 virus reads. A legend was added as a reference for size. The percentage of cbVG reads within the last 3000bp of the RSV genome (red box) are indicated in red above each plot.

**Table 2:** RNAseq/VODKA2 data of clinical samples.

| Sample | Median break position (IQR) | Median rejoin position (IQR) | Median predicted cbVG size in bp (IQR) |
| --- | --- | --- | --- |
| cl2 | 3286 (2222-5134) | 3352 (2300-5170) | 23835 (20160-25942) |
| cl59 | 3661 (2249-4935) | 3705 (2333-4982) | 23017 (20526-25828) |
| cl79 | 2981 (2000-4226) | 3030 (2032-4267) | 24445 (21969-26428) |
| cl87 | 4361 (3014-4794) | 4377 (3031-4820) | 21726 (20854-24422) |
| cl99 | 4582 (3556-4794) | 4608 (3589-4820) | 21275 (20854-23317) |
| cl103 | 3892 (3027-4794) | 3923 (3060-4821) | 22655 (20854-24370) |

IQR = interquartile range

### 3.4 Virus replication levels drive cbVG species selection

We next investigated what drives the difference in cbVG populations at passage 20 between A549 control cells and A549 STAT1 KO cells. As cbVGs are dependent on the virus replication machinery to be generated and replicated, we hypothesized that STAT1 exerts selective pressure on cbVGs by limiting virus replication and drives their evolution within a virus population. To determine how cbVG populations evolved over time in control and STAT1 KO cells, we analyzed passages 0, 4, 8, 12, 16 and 20 using our RNAseq/VODKA2 pipeline for all 3 lineages.

To assess this hypothesis, we first compared virus replication kinetics, using virus reads as a proxy, to the cbVG population diversity (Shannon Entropy) kinetics in control and STAT1 KO cells **(Figure 4A and 4B)**. We noticed that cbVG population diversity kinetics followed virus replication kinetics **(Figure 4A and 4B)**. These data also show that a permissive host environment, such as STAT1 KO cells, that allows RSV replication to be sustained at high levels will promote a more diverse cbVG population early on (passage 4, 8 and 12). As Shannon Entropy takes in consideration both richness and evenness, we also analyzed cbVG species richness kinetics and the proportion of cbVGs within the last 3000 bp of the RSV genome over time. We observed that cbVG species richness kinetics also followed virus replication kinetics and that RSV generated more cbVG species early on (passage 4, 8 and 12) in STAT1 KO cells **(Figure 4C)**. Interestingly, at passage 16 and 20, Shannon Entropy and cbVG species richness drop significantly in STAT1 KO cells and are even lower than control cells. To understand this observation, we next looked at the proportion of cbVGs within the last 3000 bp of the RSV genome (= predicted short cbVGs) **(Figure 4D)**. We measured this proportion in two ways: (1) percentage of predicted short cbVG species found in the sample (empty circles) and (2) percentage of predicted short cbVG reads found in the sample (full circles). In STAT1 KO cells, at passage 16 and 20, RSV generated more of the shorter cbVGs species (pink empty circles) that also accumulated significantly (pink full circles) **(Figure 4D)**. These data suggest that predicted shorter cbVGs outcompeted the predicted longer cbVG species, which led to the drop in cbVG population diversity and richness in STAT1 KO cells at passage 16 and passage 20. Importantly, only when the predicted short cbVGs predominated the cbVG population (passage 20), we observed a significant increase in cbVG content in STAT1 KO cells **(Figure 4E)**. In rare circumstances, such as passage 8 from control lineage 3, where virus reads dropped significantly but a few cbVGs reads could still be detected (13 cbVG reads/6073 virus reads), cbVG content was also high in relationship to virus reads **(Figure 4E)**. If this is technical (e.g. threshold of detection) or biological (e.g. predicted long cbVGs are very stable in the absence of helper virus) is unknown. We also tracked the most predominant cbVG species in the three STAT1 KO cells lineages and observed that they were all not present at P0 and all were detected first at passage 16 **(Figure 4F–4H)**. This indicates that these predicted shorter cbVGs were *de novo* generated before being selected and outcompeting over cbVGs.

**Figure 4:**
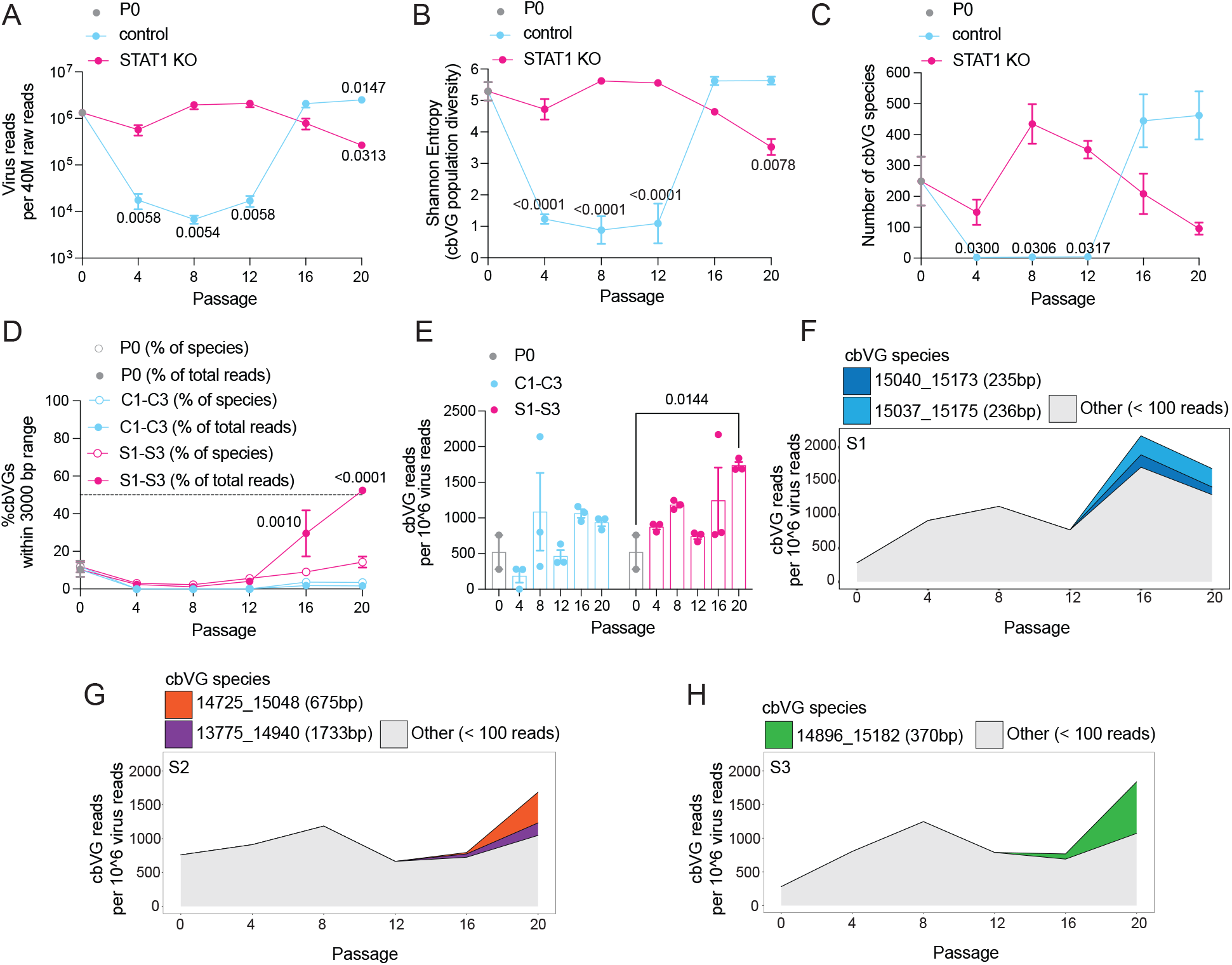
Sustained high replication levels in A549 STAT1 KO drive the selection of short cbVG species. (A) Line graph shows virus reads normalized per 40 million total raw reads for different passages in A549 control (blue) or STAT1 KO (pink) cells. (B) Line graph shows Shannon Entropy for different passages in A549 control (blue) or STAT1 KO (pink) cells. (C) Line graph shows cbVG species richness for different passages in A549 control (blue) or STAT1 KO (pink) cells. (D) Line graph shows % cbVGs species (empty circles) or reads (full circles) within the last 3000 bp of RSV genome for different passages in A549 control (blue) or STAT1 KO (pink) cells. (E) Bar graphs shows cbVG reads normalized to 1 million virus reads. (A-E) Data are shown as mean ± s.e.m. P values for two-way analysis of variance (ANOVA) with Bonferroni post-hoc test comparing all passages (n= 3 lineages) to P0 (n= 2). (F-H) Continuous stacked bar graph shows cbVG species with at least 100 reads present in each sample of lineage 1, 2 and 3 of A549 STAT1 KO. The legend includes information on break position, rejoin position and predicted cbVG size. cbVG species with less than 100 reads were grouped together as “Other”.

Overall, these data show that longer predicted cbVG species are predominant in both cell lines until sustained high virus replication levels lead to diversification of cbVGs species and generation of shorter cbVGs. In STAT1 KO cells, where high virus replication levels were maintained throughout the experiment, we observed that shorter cbVGs were selected and accumulated, which allowed for an increase in the overall cbVG content within the virus population. However, in control cells this selection was not observed, likely because high virus replication levels were not maintained throughout the experiment and peak virus replication levels were not reached before passage 20.

To more directly address if virus replication levels are the major driver of cbVG selection, we serially passaged RSV at a fixed MOI of either 0.1 or 10 in both A549 control and STAT1 KO cells for 10 passages. As opposed to the fixed-volume experiment, these conditions would normalize virus replication levels between control and STAT1 KO cells throughout the experiment. MOI of 0.1 was chosen to maintain virus replication levels low and MOI of 10 was chosen to maintain virus replication levels high. As with the fixed-volume experiment **(Figure 1A)**, we carried out three experimental evolution replicates for each cell line. We hypothesized that if sustained high replication levels are the major driver of short cbVG selection, we should observe selection of short cbVGs only at a fixed-MOI of 10 but in both cell lines due to normalization of virus replication levels.

As expected, RSV replication levels, as measured by titer, were similar between control and STAT1 KO cells throughout the experiment for both MOIs **(Figure 5A and 5B)**. We analyzed P0 and P10 viral stocks using the RNAseq/VODKA2 pipeline and found that RSV populations contained more cbVGs when passaged at an MOI of 10 than 0.1 in both cell lines **(Figure 5C)**. At MOI 10 we noticed an accumulation of cbVGs with break and rejoins occurring in the last 3000 bp of the RSV genome (red box) corresponding with cbVGs of short length regardless of the cell line **(Figure 5D)**. This did not occur at an MOI 0.1 where mainly longer cbVG were present **(Figure 5D)**. To assess if the differences observed were consistent between lineages, we plotted the median break, rejoin and predicted size values for each lineage in each condition **(Figure 5E–5G)**. All three analyses showed more drastic differences between MOI 0.1 and 10 than among control and STAT1 KO cells, which we expected due to normalization of virus replication levels between cell lines **(Figure 5E–5G)**. At MOI 10, we noticed selection of shorter cbVG species that dominated the cbVG population **(Figure 5H)**, similar to what we observed in STAT1 KO in the fixed-volume experiment **(Figure 2E)**. Notably, at passage 10, some of the most predominant cbVGs species were of longer predicted length than the most predominant species at passage 20 in the fixed-volume experiment suggesting that the average length of accumulated cbVG species get shorter over time **(Figure 2E and Figure 5H)**. Finally, we did observe that shorter cbVG species tended to accumulate more in STAT1 KO than control cells during MOI 10 passages, suggesting that other factors might play a minor role alongside virus replication levels **(Figure 5D–5H)**.

We also compared the fixed-volume versus fixed-MOI experiments more directly **(Figure S2)**. Virus titers in the fixed-volume control and MOI 0.1 experiments never went above 2×10^7 TCID_50_/mL (see dashed line), however in the fixed-volume STAT1 KO and MOI 10 experiments virus titers were sustained above this threshold over longer periods time **(Figure S2A)**. We also compared the distribution of break, rejoin and size between the two experiments and observed a shift in distribution only in the fixed-volume STAT1 KO and MOI 10 experiments **(Figure S2B–S2C)**. These data again emphasize how virus replication levels need to be sustained above a certain threshold to generate and accumulate predicted shorter cbVGs.

**Figure 5:**
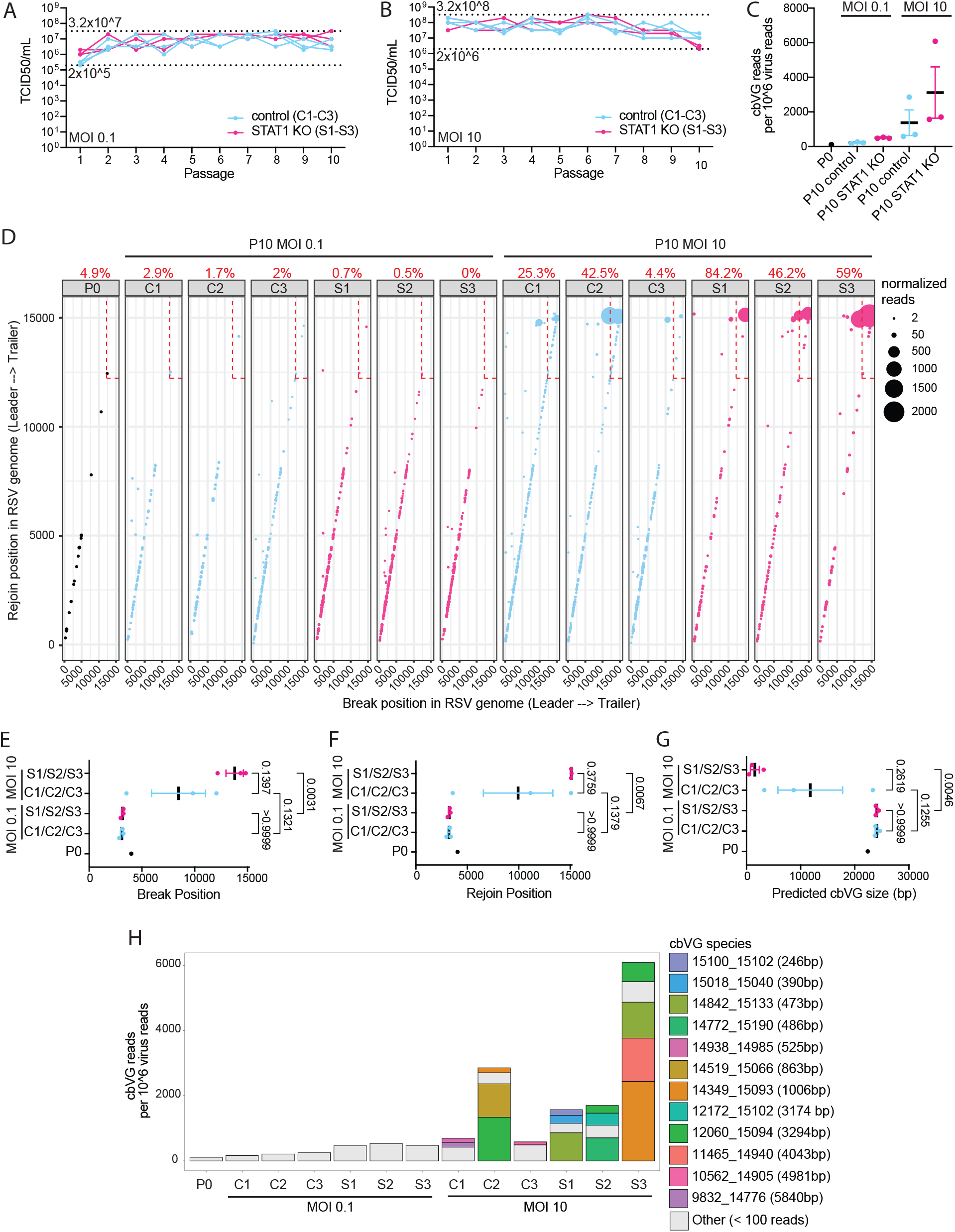
Selection of short cbVG species only occurs at high MOI conditions. (A-B) TCID_50_ assay was performed on Hep2 cells for all samples from fixed MOI 0.1 and 10 experiment. Dashed lines indicate the minimum and maximum TCID_50_/ml values reached in each experiment. (C) cbVG reads detected by VODKA2 were normalized per every 1 million virus reads. Each dot represents the median of a lineage. Data are shown as mean ± s.e.m. (D) Each dot represents a cbVG species and its break/rejoin position. The size of the dots represents the number of normalized reads per 10^6 virus reads. A legend was added as a reference for size. The percentage of cbVG reads within the last 3000bp of the RSV genome (red box) are indicated in red above each plot. (E-G) Mean break position, rejoin position and predicted cbVG size are shown in these graphs. Each dot represents the median of a lineage. Data are shown as mean ± s.e.m. P values for one-way analysis of variance (ANOVA) with Bonferroni post-hoc test for all groups are indicated. (H) Stacked bar graph shows cbVG species with at least 100 reads present in each sample. The legend includes information on break position, rejoin position and predicted cbVG size. cbVG species with less than 100 reads were grouped together as “Other”.

Taken together, these data confirm that sustained high MOI (high replication) conditions are a main driver of the generation and selection of short cbVGs that accumulate within virus populations. We also show that more resistant cell lines, such as A549 control, which were not susceptible to cbVG accumulation in the fixed-volume experiment, can change phenotype if saturated by high MOI conditions. Meanwhile, the more permissive cell line A549 STAT1 KO, which was susceptible to cbVG accumulation in the fixed-volume experiment, can change phenotype if MOI conditions are maintained to very low levels.

## 4. Discussion

Although non-standard viral genomes play a crucial role in shaping virus infection outcomes (Felt et al., 2021; Sun et al., 2015; Vasilijevic et al., 2017), the principles that drive the selection and evolution of non-standard viral genomes within a virus population are poorly understood. *In vitro*, non-standard viral genomes usually accumulate within virus populations when RNA viruses are serially passaged at a high multiplicity of infection (MOI) (Huang, 1973; Kingsbury et al., 1970; Stampfer et al., 1971; Sun and Lopez, 2016; Thompson and Yin, 2010; Von Magnus, 1954; von, 1951; Welch et al., 2020). These studies suggest that accumulation of non-standard viral genomes is favored by increasing the number of cells infected, the virus replication within cells, and/or the number of co-infected cells with standard and non-standard viral genomes. However, how non-standard viral genome populations change when an increase in the cbVG content of a virus population occurs remains unknown.

Non-standard viral genomes of the copy-back type are historically described as significantly shorter than standard viral genomes (Huang, 1973; Huang and Wagner, 1966; Kingsbury et al., 1970; Lazzarini et al., 1981; Vignuzzi and Lopez, 2019) and it is unclear if the bias for a shorter size is due to preferential generation of short cbVGs, faster replication rate and accumulation of short cbVGs, and/or due to limitations in detection methods. It is also not clear how cbVG species are selected for under different environmental conditions and how this selection impacts the evolution of the overall cbVG population. In this study, we address how cbVG selection and cbVG population evolution is affected by a permissive host environment that allows sustained high replication levels (A549 STAT1 KO cells) and a resistant host environment that limits replication levels (A549 control cells). Overall, we found that cbVGs of a longer predicted size predominate first and that sustained high RSV replication levels lead to cbVG population diversification. This diversification primes the generation and selection of shorter cbVGs that later accumulate and allow for an increase in the cbVG content within a virus population **(Figure 7)**.

It is important to note that although permissive STAT1 KO cells accumulated significantly more cbVGs than resistant control cells, cbVGs were found in both cell lines **(Figure 1B and Table 1).** These data suggest that accumulation but not generation of cbVGs is dependent on sustained high virus replication. This is consistent with our recent study where we showed that cbVGs were found in nasal washes of experimentally RSV-infected adult patients before high virus replication levels were reached (Felt et al., 2021). Taken together, our study also shows that generation of cbVGs is inherent to RSV replication and that most RSV populations will contain cbVGs. This is likely true for many other RNA viruses and, as cbVGs have been shown to play a crucial role in virus infections, underlines the need to consider cbVGs in the study of RNA virus populations.

Next-generation sequencing and pipelines to analyze virus populations have improved significantly in recent years (Beauclair et al., 2018; Bosma et al., 2019; Boussier et al., 2020; Jaworski and Routh, 2017; Olmo-Uceda, 2022; Routh and Johnson, 2014; Timm et al., 2014). In this study, we used an updated version of our previously published RNAseq/VODKA pipeline, called VODKA2, to study cbVG populations (Sun et al., 2019). Previous methods such as RT-PCR more efficiently detect short cbVGs that are of high abundance, however, our RNAseq/VODKA2 pipeline can overcome these challenges and detect unbiasedly cbVG species of various predicted sizes and abundance **(Figure 2A)**. Therefore, VODKA2 revealed that the diversity of cbVGs generated by RSV is much larger than previously thought. The use of stranded total RNA library preparation kits can further allow VODKA2 to determine if cbVGs are only generated (only genomic) or are also being replicated by the viral polymerase (mixture of genomic and antigenomic). From a stranded library preparation on P0, P20 C1-C3 and P20 S1-S3, we were able to determine that 39.77% of cbVGs were genomic and 60.23% of cbVGs were antigenomic indicating that cbVGs were not only generated but also replicated **(Figure S3)**. Furthermore, short cbVGs (especially shorter than 6000 bp) correlated with more reads indicating that short cbVGs are more likely to replicate more efficiently as has been shown previously for some non-standard viral genomes **(Figure S4)** (Laske et al., 2016; Mendes and Russell, 2021; Pelz et al., 2021). Overall, these data suggest that sustained high replication drives the generation of shorter cbVGs; however, it is likely the highly efficient replication of short cbVGs that ultimately leads to their accumulation and an overall increase in the cbVG content within a virus population.

Our RNAseq/VODKA2 pipeline also allowed us to compare cbVG species between virus populations from the cell line that limited virus replication (A549 control cells) and the cell line that allowed high virus replication (A549 STAT1 KO cells). The virus population observed in STAT1 KO cells mainly consisted of shorter cbVGs, whereas virus population from control cells consisted mainly of much longer cbVGs. A fixed-MOI passaging experiment confirmed that it is sustained high virus replication levels that drive the selection of shorter versus longer cbVGs. In the fixed-volume passaging experiment, we observed that sustained high virus replication over several passages drives cbVG diversification and generation of short cbVGs that will significantly accumulate **(Figure 4)**. This demonstrates that high virus replication does not directly lead to cbVG accumulation and instead requires an intermediate step that consists of cbVG diversification and generation of short cbVGs, which then accumulate and take over the cbVG population **(Figure 6)**. The exact molecular mechanism that drives the generation of shorter cbVGs is not clear. Are shorter cbVGs generated from longer ones? Do high virus replication levels affect the processivity of the polymerase directly or maybe indirectly through host factors? Future studies will address these questions.

**Figure 6:**
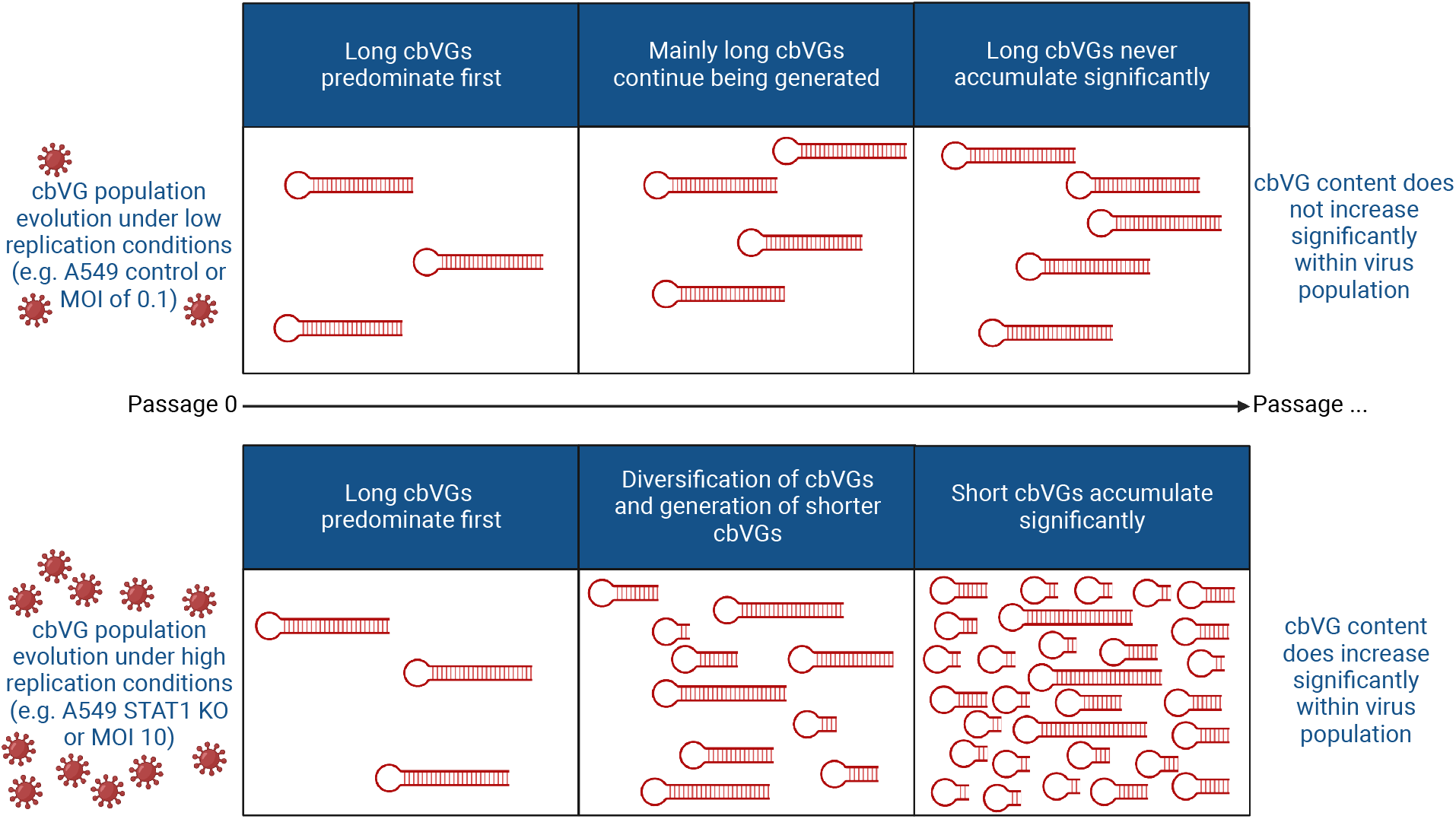
Schematic summary of cbVG population evolution in resistant and permissive host. Summary of the cbVG population changes observed in our study over time in different host environments.

In this study, we also discovered the first evidence of cbVGs of long predicted sizes. These cbVGs are predominant at first *in vitro* and are the most predominant *in vivo*. Some of these cbVGs are predicted to be longer than the genome **(Figure S1B)**. It is possible that previous studies did not detect these longer cbVGs due to lack of sensitivity in the methods used. The fact that we detected these longer cbVGs in patients raises the question on how they are functionally different to the shorter cbVGs, which have been studied so far *in vitro*. Do they interact with different pattern recognition receptors and/or induce different signaling pathways? Why do we mainly find longer cbVGs in patients and less shorter ones? Is there a fitness advantage of these longer cbVGs *in vivo*? We must also consider that we are limited to nasal samples from patients and that cbVG populations could differ between microenvironments (e.g. nose vs lung) as has been shown for Zika non-standard viral genomes (Johnson et al., 2020). However, we have observed that Sendai virus (SeV) populations in the lung of mice also consist of mainly longer cbVGs, indicating that this might be a common feature of *in vivo* cbVG populations (unpublished data). Further characterization of longer cbVGs is still needed as we can only extract predicted sizes based on VODKA2. Due to their size and complex structure, amplifying or sequencing the entire sequence of long cbVGs by RT-PCR or MinION remains a challenge. So far, we were able to confirm that the junction regions at least are not artifactual as we did not detect copy-backs when running VODKA2 using a RSV database (negative control for false positivity rate of VODKA2) or GAPDH/actin databases (negative controls for library prep generating copy-back sequences) on SeV samples **(Figure S5A)**. Interestingly, cbVG species of longer predicted size never accumulated to significant amounts, strongly suggesting that they are indeed of longer size and hence not as competitive against the standard viral genome or shorter cbVG species **(Figure S4)**. However, it is important to note, that many predicted long cbVG species can be detected in several passages and follow virus kinetics, which suggests that they could potentially be replicated and packaged **(Figure S5B)**.

Non-standard viral genomes can interfere with standard viral genome replication when they accumulate to significant levels causing a significant drop in infectious virus particles (Genoyer and Lopez, 2019; Kirkwood and Bangham, 1994; Manzoni and Lopez, 2018; Palma and Huang, 1974). Although cbVGs did accumulate to significant levels in STAT1 KO cells, we did not observe such a significant decrease in infectious virus particles in the fixed-volume experiment **(Figure 1B)**. This is likely due to cbVGs being less interfering in the absence of antiviral responses (e.g. mediated by STAT1), such as has been shown for RSV and parainfluenza virus 5 (Sun et al., 2015; Wignall-Fleming et al., 2020).

Currently, they are few studies showing what roles host proteins could play in non-standard viral genome generation, accumulation, and selection (Holland et al., 1976; Kang and Allen, 1978). We have previously shown that children are more likely to accumulate RSV cbVGs than adults and that female patients are more likely to accumulate RSV cbVGs than males (Felt et al., 2021). Furthermore, in an adult cohort, although all patients were experimentally infected with the same virus (RSV Memphis 37), not all patients accumulated cbVGs (Felt et al., 2021). All these studies strongly suggest that the host might play an important role in shaping cbVG populations. In this study, we have shown that STAT1 by limiting virus replication can exert selective pressure on cbVG populations and shape their evolution. It will be important to assess in future studies what other host proteins can shape cbVG content within virus populations.

## Acknowledgments

We thank Dr. Susan Weiss for kindly providing A549 control and STAT1 KO cells. Figures 1A, S1A and 6 were created with BioRender.com.

## Funding

This works was supported by the US National Institutes of Health National Institute of Allergy and Infectious Diseases (AI137062 and AI134862 to CLB).

## Conflict of interest

None declared.

## Data Availability

Raw sequence data are deposited on SRA (PRJNA837014).

## Supplementary Data

**Figure S1:**
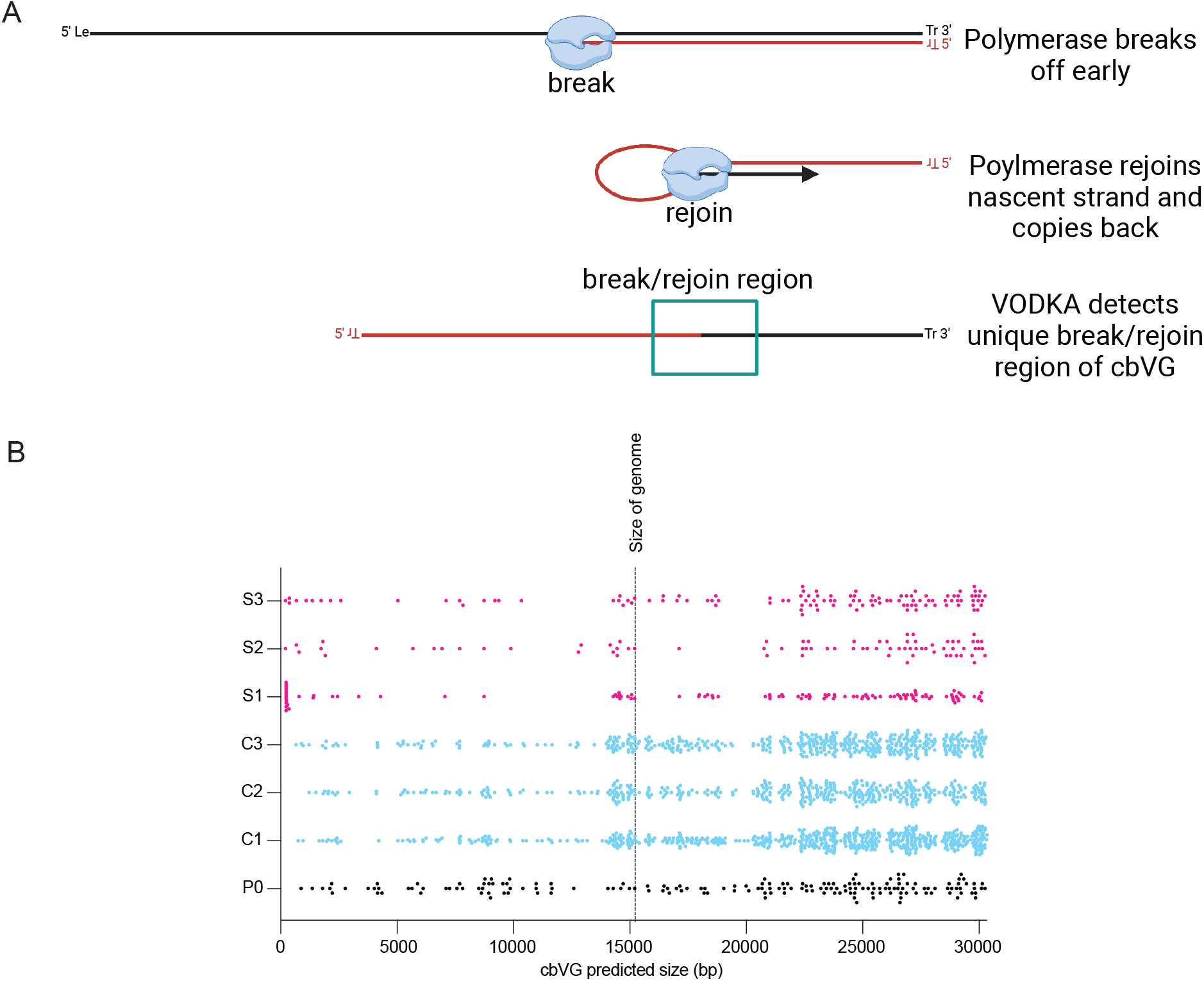
RNAseq/VODKA2 approach and distribution of predicted cbVG sizes. (A) Schematic of cbVG generation and cbVG detection by VODKA2. (B) Data represents cbVG predicted size distribution for P0, P20 C1-C3 and P20 S1-S3. Each dot represents a cbVG species. The black dashed line indicates the size of the genome (15223bp).

**Figure S2:**
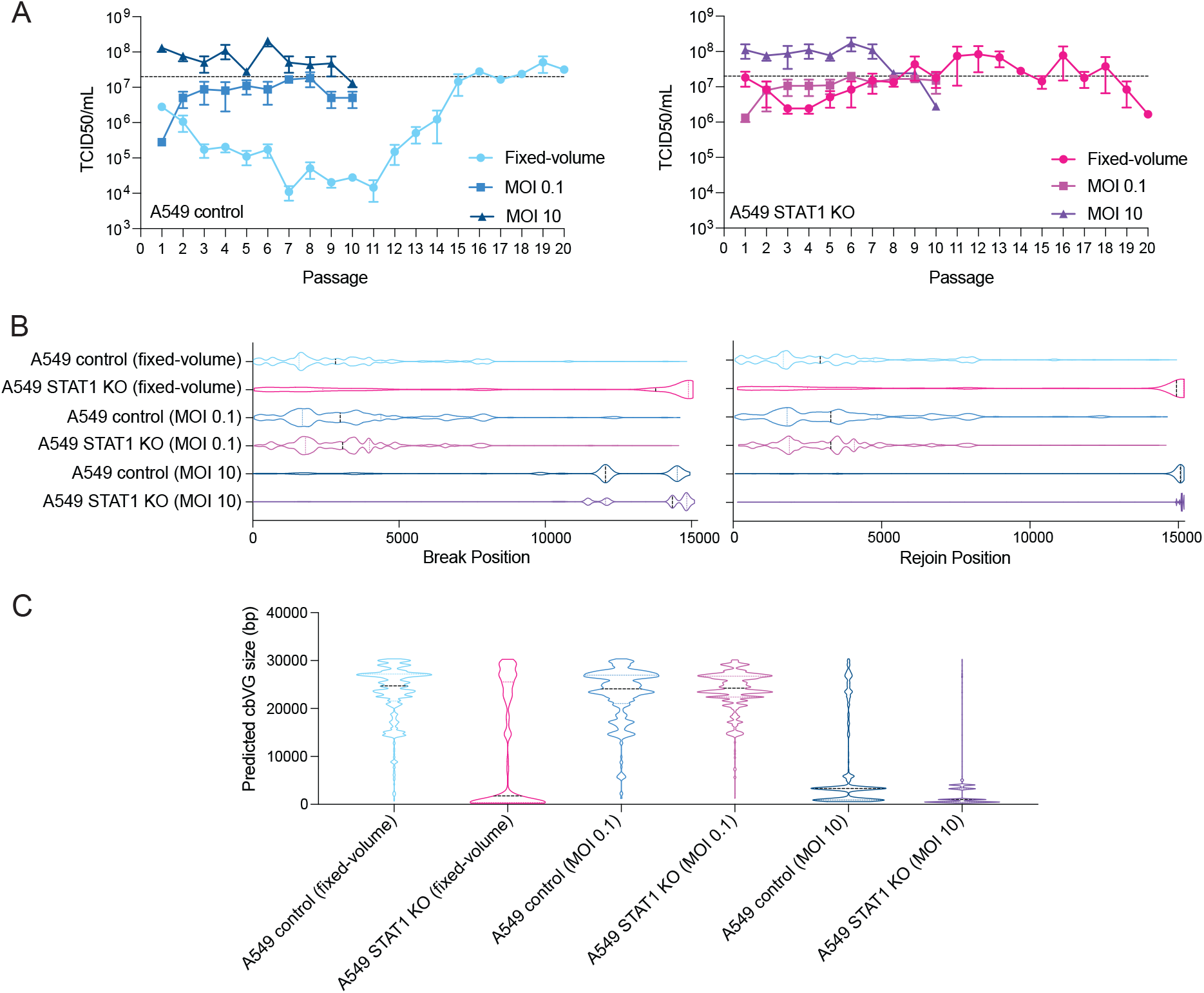
Comparison of virus kinetics and cbVG populations between fixed-volume and fixed-MOI experiments. (A) TCID_50_ assay was performed on Hep2 cells for all samples (n= 3 lineages). Dashed lines indicate 2×10^7 TCID_50_/ml, which is highest titer MOI 0.1 reached. (B) Graphs represent distribution of break and rejoin positions. Data are shown as truncated violin plots with median and interquartile ranges (n= 3 lineages). (C) Graphs represent distribution of predicted sizes of cbVGs. Data are shown as truncated violin plots with median and interquartile ranges (n= 3 lineages).

**Figure S3:**
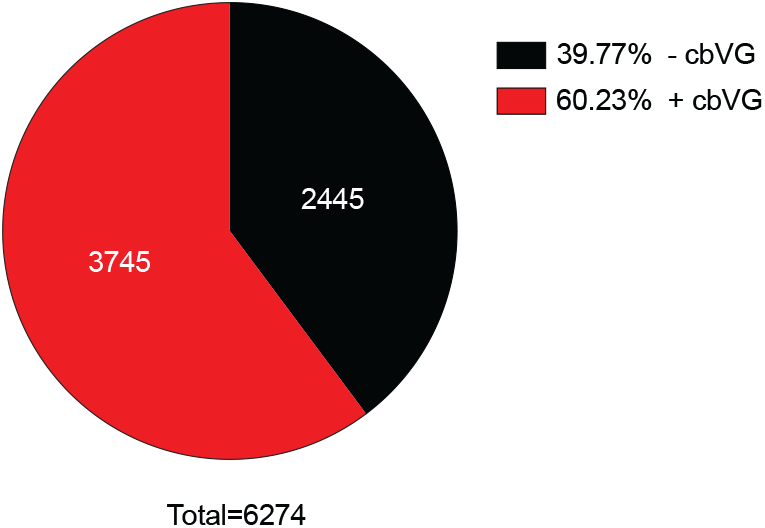
Proportions of genomic and antigenomic cbVGs. cbVGs from fixed-volume experiment (P0, P20 C1-C3 and P20 S1-S3) were aligned to reference genome to assess strandness. Pie chart indicates proportion of genomic (-cbVG) and antigenomic (+cbVG) cbVGs.

**Figure S4:**
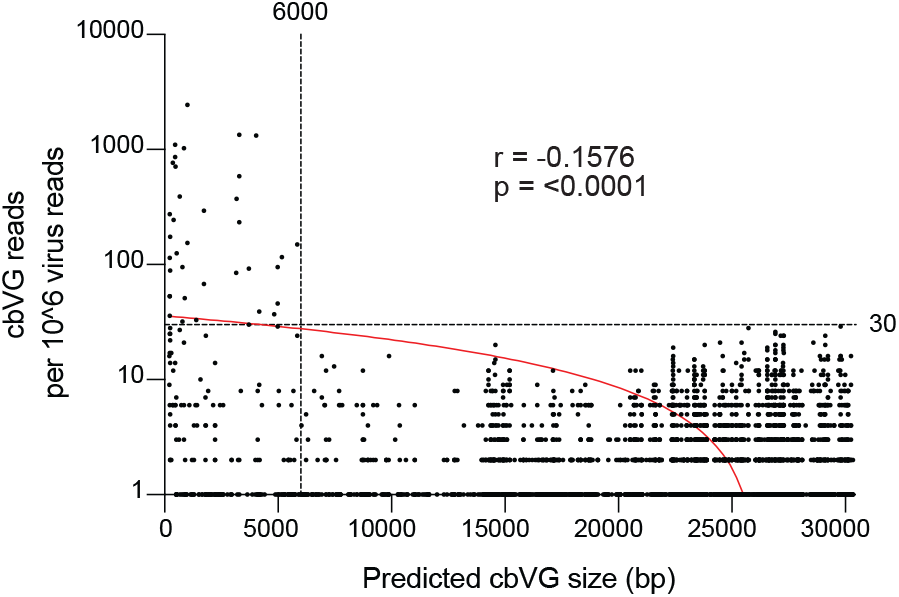
Correlation between predicted cbVG size and cbVG reads. Each dot represents a cbVG species from P20 fixed-volume experiment (C1-C3 and S1-S3) and from P10 fixed-MOI 0.1 and 10 experiments (C1-C3 and S1-S3). P value is shown for Pearson correlation between predicted cbVG size and cbVG reads are indicated. Line of best fit for simple linear regression is shown in red. Black dashed lines indicate 30 reads and 6000 bp.

**Figure S5:**
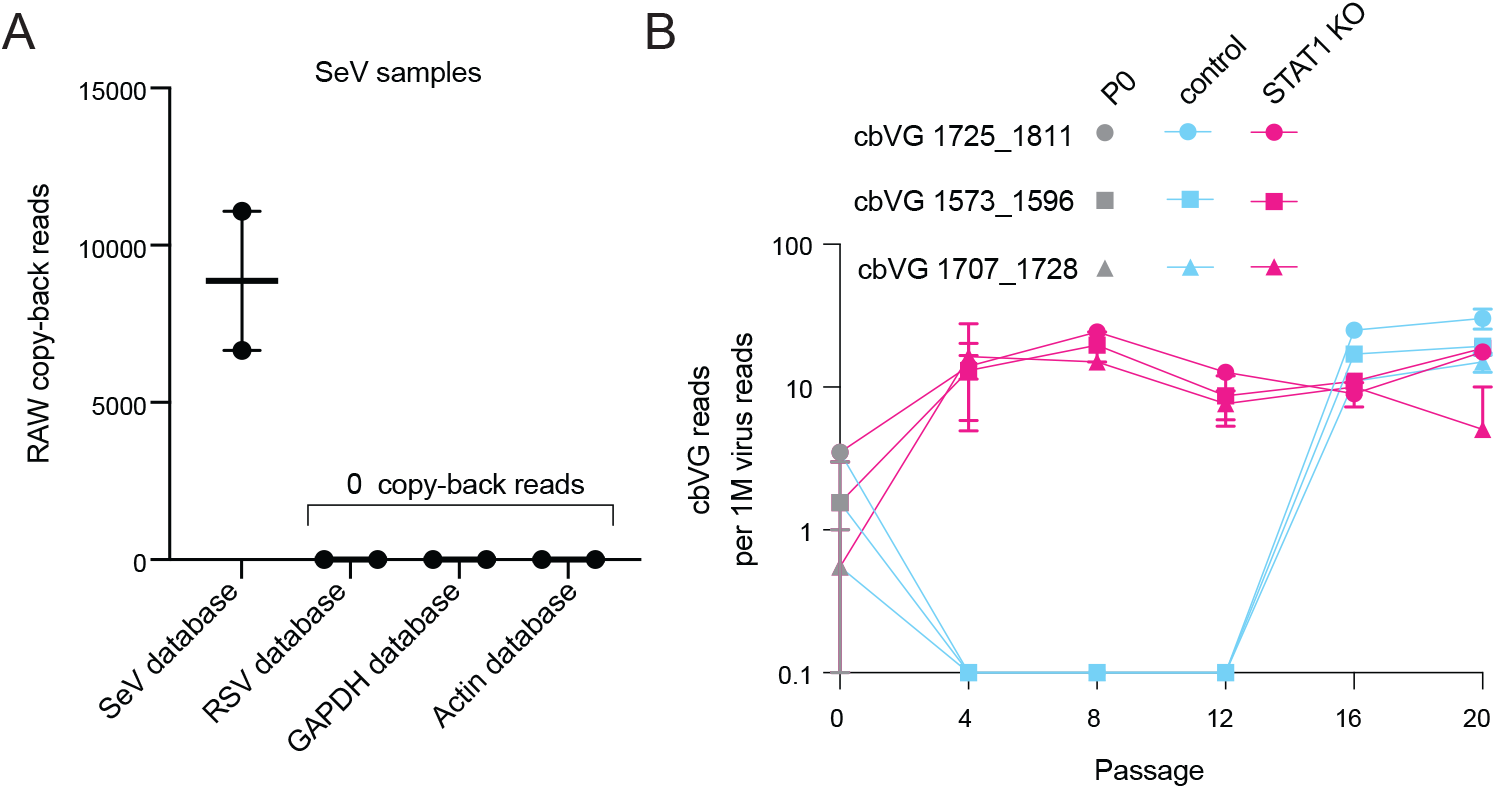
Confirmation and tracking of predicted long cbVG species. (A) Two SeV samples from a previous study (Sun et al., 2019; Xu et al., 2017) were analyzed by VODKA2 with an SeV, RSV, GAPDH or Actin database. Raw data are shown as mean ± s.e.m. (n= 2) (B) Line graph shows number of reads for 3 different cbVG species normalized per 1 million virus reads for different passages in A549 control (blue) or STAT1 KO (pink) cells (n= 3 lineages). The break and rejoin positions are indicated in the legend.

## Notes

### Competing Interest Statement

The authors have declared no competing interest.

### Summary of Updates

Specifically, we changed: Figure 1 and Figure 2 now contain data with same library prep and sequencing methodology than Figure 4 and 5. Figure 4 now contains data for all 3 lineages.

